# A spatiotemporal immune atlas of subarachnoid hemorrhage from single-cell and spatial transcriptomics

**DOI:** 10.64898/2026.02.02.703421

**Authors:** Changming Liu, Bingrui Zhu, Yuchun Liu, Qian Yu, Yongkang Yi, Jinchen Zhou, Xiaocan Wang, Canyu Ma, Yibo Liu, Guangmin Qiu, Haifeng Chu, Kaikai Wang, Jianmin Zhang, Xiaoyu Wang

## Abstract

**Background and Purpose:** Subarachnoid hemorrhage (SAH) triggers a complex immune response that critically influences early brain injury (EBI) and long-term outcomes. However, the precise spatiotemporal dynamics and heterogeneity of immune cell infiltration and microglial reprogramming remain poorly understood. We aimed to construct a high-resolution immune atlas to delineate cell states, lineage trajectories, and spatial niches following SAH.

**Methods:** We integrated single-cell RNA sequencing (scRNA-seq) of CD45+ immune cells with spatial transcriptomics (ST) in a murine endovascular perforation SAH model. Immune landscapes were profiled at 24 hours (acute phase) and 72 hours (subacute phase) post-injury, compared with sham controls. Advanced bioinformatics integrated transcriptional signatures with spatial localization to map macrophage, neutrophil, and microglial dynamics.

**Results:** Our atlas reveals a coordinated immune transition from acute inflammation to reparative processing. We identified five macrophage, four neutrophil, and eight microglial subsets with distinct spatiotemporal patterns. Notably, we discovered a SAH-specific inflammatory microglial population (MG_03; *Spp1+*/*Lpl+*) that clusters at the rupture site during the acute phase. This subset is transcriptionally distinct from disease-associated microglia (DAM) in other neurodegenerative conditions. Trajectory analysis suggests MG_03 acts as a signaling hub for immune recruitment before transitioning toward proliferative and reparative states (MG_06–08) that disperse into the parenchyma by 72 hours.

**Conclusions:** This study provides the first comprehensive spatiotemporal immune atlas of SAH, highlighting the distinct role of the *Spp1+* MG_03 subpopulation in early injury sensing. These findings offer a roadmap for identifying precise therapeutic windows and targeting specific immune subsets to mitigate EBI.

**Graphical Abstract:** Experimental workflow for single-cell RNA sequencing (scRNA-seq) and spatial transcriptomics (ST) in a mouse SAH model induced by endovascular perforation. Brain tissue from the ipsilateral (injured) hemisphere was collected from sham and at 24 h and 72 h post-SAH. For scRNA-seq, CD45⁺ immune cells were isolated prior to library preparation.

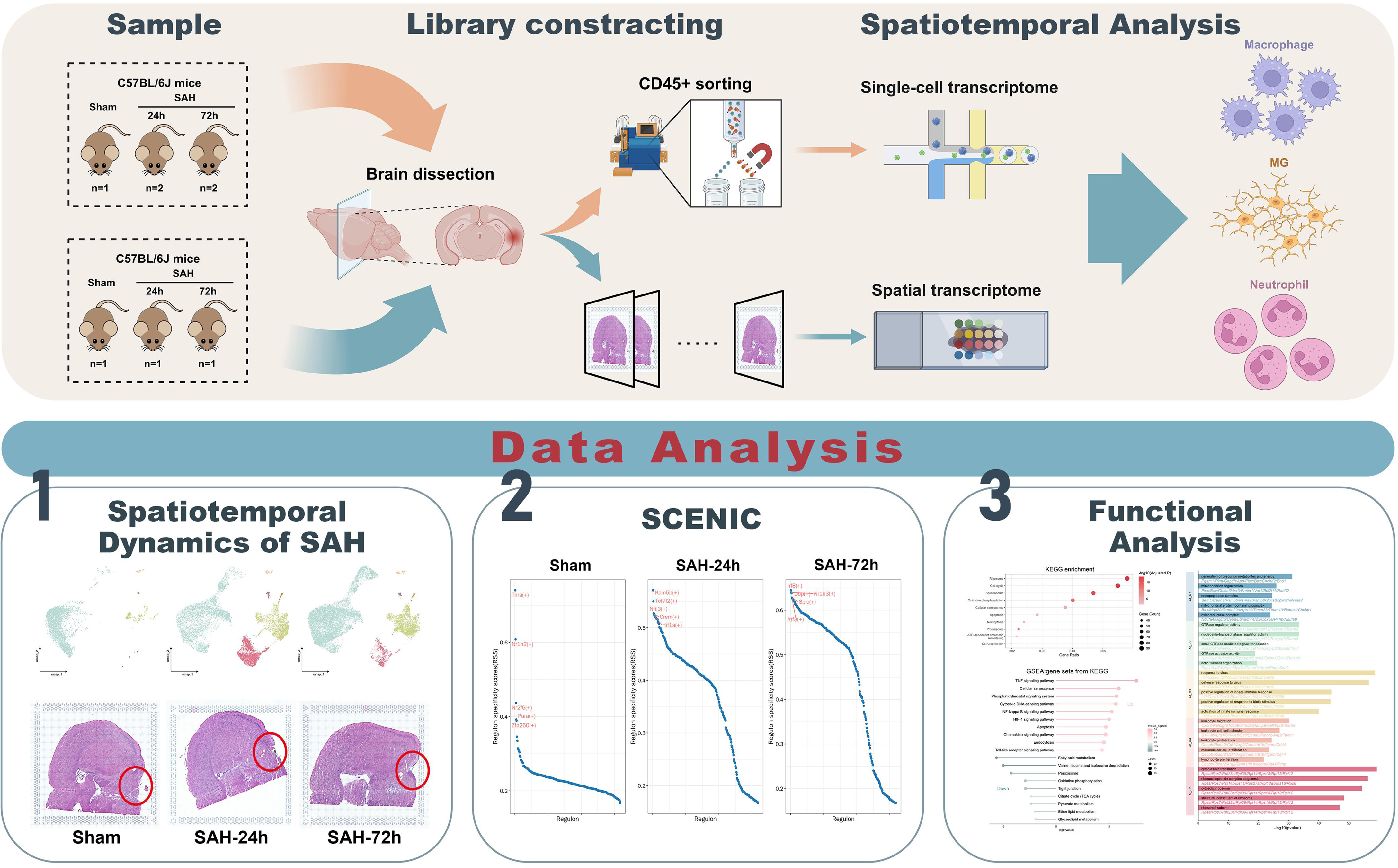

## Introduction

Neuroimmunity has emerged as a major frontier in neuroscience, shaping both normal brain function and disease pathogenesis across the lifespan^1^. Immune signaling contributes to learning and memory^1^, neural development, stress responses, and repair^2^, whereas dysregulated responses are implicated across a broad spectrum of central nervous system (CNS) disorders^3^. In neurodegenerative diseases such as Alzheimer’s disease and Parkinson’s disease, persistent neuroinflammation is thought to accelerate pathological protein deposition and neuronal loss^4^. In multiple sclerosis, maladaptive immunity drives demyelination within the CNS. Following stroke and traumatic brain injury (TBI), peripheral immune cell infiltration and excessive activation often exacerbate early damage, whereas later phases can facilitate repair. In brain tumors, the immune system retains anti-tumor potential, yet malignant cells frequently co-opt immune evasion mechanisms to promote progression^5^.

Innate and adaptive immune cells are key effectors of these processes^6^, and their dynamic states directly influence tissue injury, repair efficiency, and functional recovery. Microglia, the resident immune cells of the CNS, rapidly activate after injury and secrete pro-inflammatory mediators such as IL-1β and TNF-α to clear damaged tissue^7^; during the repair phase, they can adopt anti-inflammatory phenotype that support neuroprotection and tissue regeneration. Peripherally derived macrophages infiltrate early, initially amplifying inflammation before transitioning to reparative states that aid efferocytosis and tissue remodeling. Lymphocytes also shape neuroimmunity: effector T cells can intensify inflammation via IFN-γ^8^, whereas regulatory T cells (Tregs) restrain excessive immune activation through IL-10 and related mediators; B cells contribute to neuroprotection in select contexts through antibody production and modulation of innate immune cells^9^.

Despite these advances, the pronounced spatiotemporal heterogeneity of neuroimmune responses remain difficult to resolve with traditional bulk RNA sequencing or histological approaches ^10^. Recent advances in single-cell RNA sequencing (scRNA-seq) and spatial transcriptomics (ST) now enable joint dissection of cell states, anatomical niches, and temporal trajectories at high resolution, providing an integrated framework to decode neuroimmune dynamics in vivo. ^,11^.

Subarachnoid hemorrhage (SAH) is a neurosurgical emergency with high early mortality and long-term disability^12^. Early brain injury (EBI), occurring within the first 72 hours, is a major determinant of clinical outcome, yet how innate immune cells are reorganized across space and time after SAH remains incompletely defined. Although prior studies have highlighted the importance of immune cells contributions to SAH, systematic, single-cell-resolved investigations that are integrated with spatial information are scarce.

To address this gap, we combined scRNA-seq of CD45^+^ cells with ST in a murine endovascular perforation model to construct a spatiotemporal atlas of immune responses at 24 and 72 hours after SAH. Our analysis delineates transcriptional states, spatial distributions, and lineage relationships of major immune populations-including macrophages, neutrophils, and microglia, and identifies SAH-associated microglial subpopulations that emerge during disease progression. These results provide a comprehensive immune landscape of SAH and offer mechanistic insights into EBI, laying groundwork for future immune-targeted therapeutic strategies.

## Results

### Spatial features of the immune-inflammatory response in SAH

We profiled spatial gene expression in a murine endovascular perforation model of SAH, which closely recapitulates clinical aneurysmal rupture^13^. At 24 h and 72 h after SAH (n = 1 per time point for spatial transcriptomic), tissue section from ipsilateral cerebral hemisphere were subjected to spatial transcriptomic profiling.

To map inflammatory activity, we computed gene-set scores (AddModuleScore) using curated inflammation-related programs (GO/Hallmark; e.g., *acute inflammatory response, positive regulation of acute inflammatory response, immune response, activation of immune response, neuroinflammatory response, and positive regulation of neuroinflammatory response*). These scores revealed widespread activation across the injured hemisphere with superimposed hotspots centered on the puncture tract and extending into the adjacent parenchyma (Fig. 1a). Consistently, immune cell-oriented gene sets indicated that macrophage activation and chemotaxis were broadly distributed within the parenchyma at 24 h but became more spatially concentrated near the puncture tract by 72 h (Fig. 1b). Leukocyte-related gene sets (leukocyte activation, migration, chemotaxis, and adhesion) were likewise enriched around the puncture site, with stronger signals at 72 h (Fig. 1c). Regions with high cell activation and microglial activation^14^ scores were centered on the puncture site and contiguous parenchyma (Fig. 1d).

**Figure 1.**
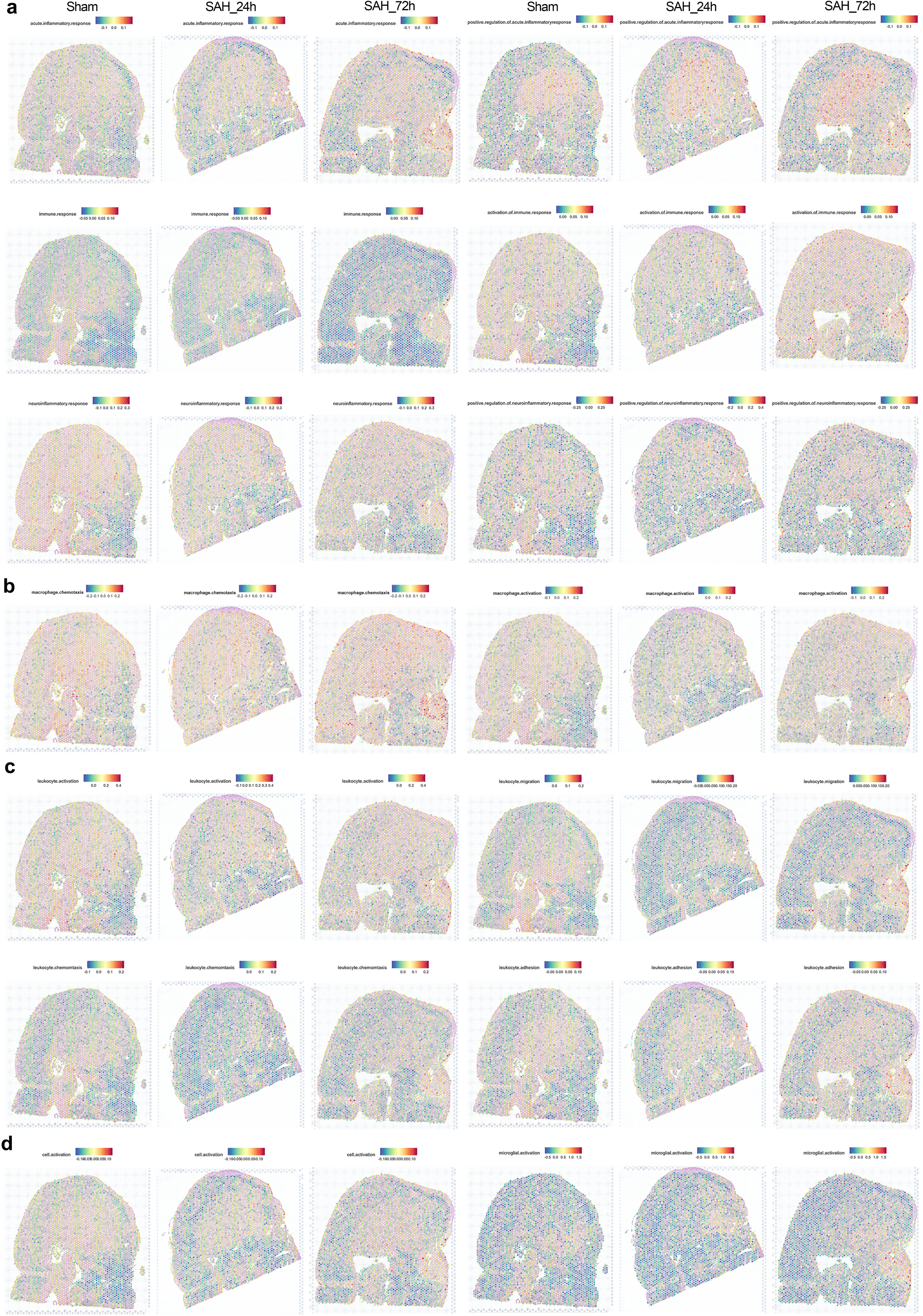
Spatial module scoring of immune programs in a mouse model of SAH. a-d, ST AddModuleScore maps for curated immune/inflammation gene sets (GO/Hallmark), illustrating pathway activity across the injured hemisphere with superimposed hotspots at the puncture tract and adjacent parenchyma. Representative programs include acute inflammatory response, activation of immune response, neuroinflammatory response, macrophage activation/chemotaxis, leukocyte activation/migration/chemotaxis/adhesion, cell activation, and microglial activation.

### Spatiotemporal dynamics of immune cells after SAH

To dissect the heterogeneity of immune cells during SAH, we isolated CD45⁺ immune cells from the ipsilateral hemisphere of mice at staged time points after SAH and performed scRNA-seq (Sham, n = 1; SAH_24h, n = 2; SAH_72h, n = 2). After quality control and filtering, 32,368 cells were retained, including 3,750 cells from sham, 10,986 from SAH_24h, and 17,632 from SAH_72h. Using an unsupervised graph-based clustering algorithm implemented in Seurat v5, we identified six major immune cell types based on canonical markers: microglia (MG), macrophage, neutrophil, T & NK cell, dendritic cell (DC), and B cell (Fig. 2a,c). Analysis of their relative abundance revealed that macrophages, neutrophils, and MG constituted the dominant populations and exhibited dynamic changes over the disease course, whereas lymphoid subsets displayed relatively modest alterations (Fig. 2b).Gene Ontology (GO) enrichment analysis corroborated these dynamics: sham samples were enriched for fundamental processes (RNA splicing, gliogenesis, and DNA repair), whereas 24 h showed strong enrichment for neutrophil migration, activation, and degranulation, consistent with acute innate responses; 72 h shifted toward antigen processing, presentation and lymphocyte-mediated cytotoxicity, indicating a transition from innate amplification to adaptive engagement (Fig. 2d). To prioritize contributors, we applied AddModuleScore for functional scoring. Macrophage, MG, and neutrophil displayed high scores for leukocyte chemotaxis, neuroinflammatory response, and positive regulation of neuroinflammatory response (Fig. 2e).

**Figure 2.**
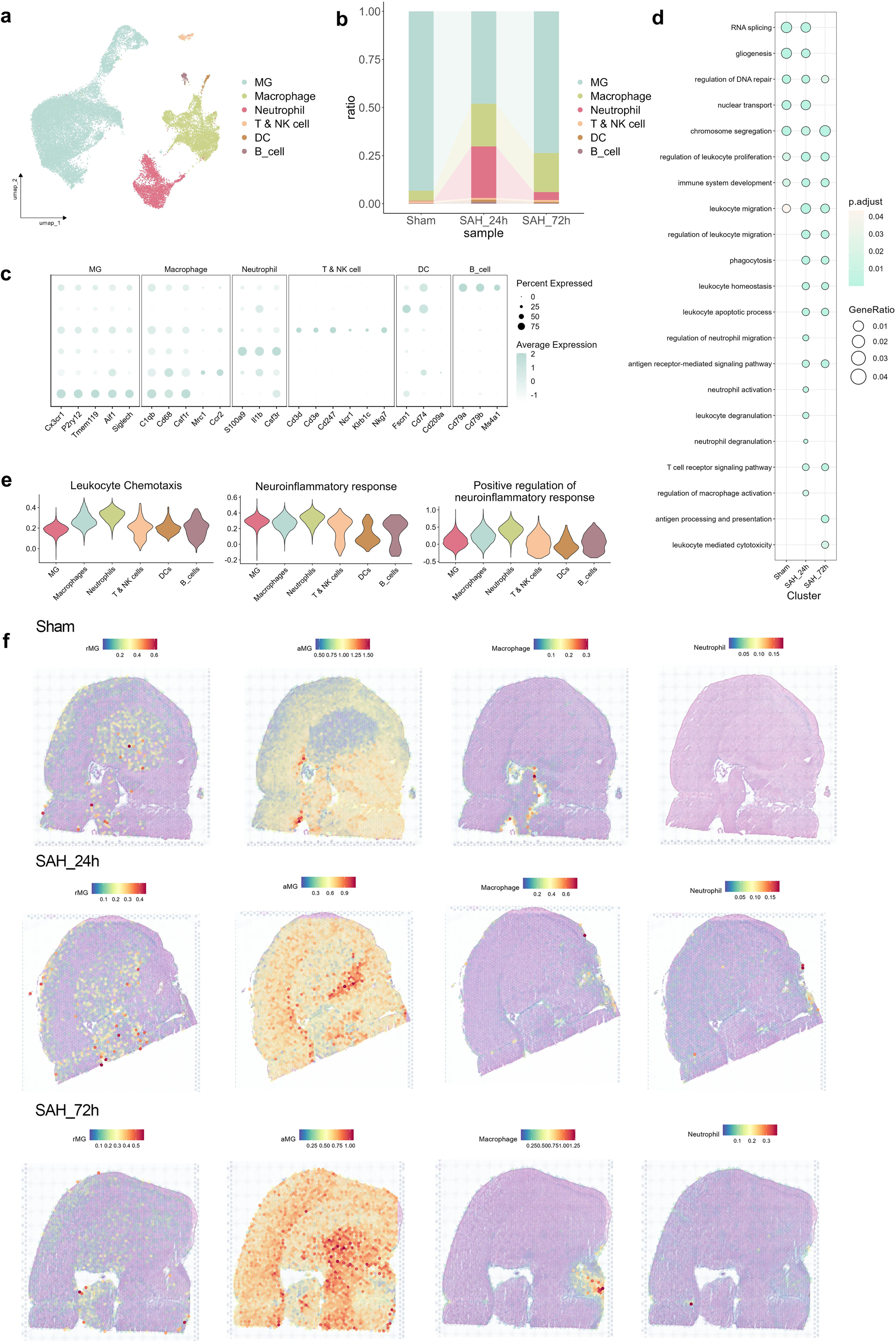
Single-cell and spatial profiling of immune populations across SAH time points. a, UMAP of 32,368 CD45⁺ cells annotated into six lineages by canonical markers: microglia (MG), macrophage, neutrophil, T & NK cell, dendritic cell (DC), and B cell. b, Relative composition of the six lineages across stages (sham, 24 h, 72 h). c, Bubble plot of representative markers per lineage/subcluster. d, Gene Ontology (GO) enrichment of DEGs across stages (BH-adjusted *P* < 0.05). e, AddModuleScore module activities for immune/inflammation programs across lineages and subclusters. f, Spatial deconvolution (RCTD) showing estimated abundances of activated microglia (aMG; P2ry12⁻), resident microglia (rMG; P2ry12⁺), neutrophils, and macrophages in sham, SAH_24 h, and SAH_72 h sections; color intensity reflects relative abundance.

Finally, we integrated scRNA-seq with ST to reconstruct the spatiotemporal distribution of immune populations. For greater specificity, MG were partitioned into resting MG (rMG; *P2ry12⁺*) and activated MG (aMG; *P2ry12⁻*). Deconvolution and low-dimensional embedding indicated rMG in diffuse parenchyma and periventricular regions, whereas aMG were sparse in sham but expanded robustly after SAH, forming dense cluster within parenchyma that intensified by 72 h. Macrophages, localized to ventricular boundaries in sham and accumulated at the puncture tract after SAH, mirroring the hemorrhage location. Neutrophils, absent as residents, concentrated at the puncture site after SAH but declined by 72 h (Fig. 2f).

### Heterogeneity of macrophages at different stages after SAH

As critical regulators of neuroinflammation, macrophages may shape the pathological process of SAH^15^. Clustering of 4,862 macrophages resolved five subpopulations with distinct signatures (Fig. 3a,b). M_01 (*Spp1⁺, Lgals1⁺*) was enriched in oxidoreductase complex and mitochondrial pathways, and likely represents a transitional functional state. M_02 (*Cd163⁺, Lyve1⁺)* aligned with border-associated macrophages, enriched in GTPase and energy metabolism, serving as a homeostatic reference. M_03 (*Ifit3⁺, Irf7⁺*) predominated at 72 h, enriched in antiviral responses and positive regulation of inflammation, representing the main effector subset of 72h-stage inflammatory injury. M_04 (*Arg2⁺, Csf3r⁺*) expanded transiently at 24 h, functionally enriched in leukocyte chemotaxis, adhesion, and proliferation pathways, defining the core subset of the acute inflammatory phase. M_05 (*H2-Aa⁺, H2-Ab1*⁺) showed ribosomal and antigen-presentation pathways enrichment, indicating strong antigen-processing potential.

**Figure 3.**
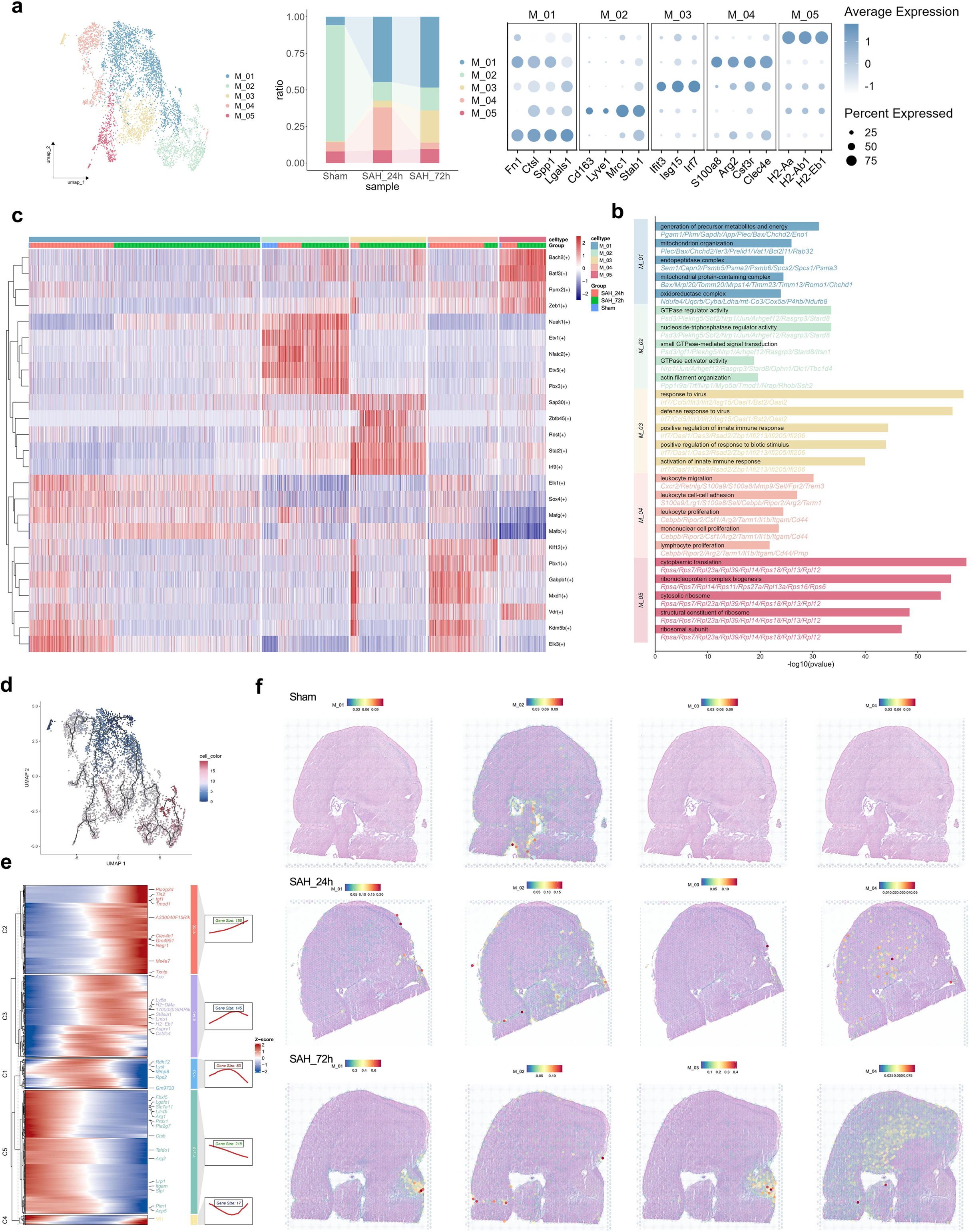
Macrophage heterogeneity and dynamics after SAH. a, UMAP resolving five macrophage subclusters (M_01–M_05); Bubble plot of representative markers per subcluster; Subcluster abundances across conditions (sham, 24 h, 72 h). b, GO enrichment of subcluster DEGs (BH-adjusted *P* < 0.05). c, Heatmap of transcription-factor activity across subclusters (top five TFs per subcluster). d, Pseudotime trajectories inferred from scRNA-seq (Monocle3/CytoTRACE2/Slingshot-concordant). e, Branch-dependent gene dynamics along pseudotime; Z-scored expression grouped into five modules with representative genes indicated. f, Spatial deconvolution maps of macrophage subclusters in sham, SAH-24 h, and SAH-72 h sections; color intensity reflects relative abundance..

To further resolve macrophage dynamics after SAH, we performed pseudotime trajectory analysis using Monocle3. Trajectory inference positioned M_01 at the root, bifurcating toward M_04 to M_05 and M_02 to M_03, consistent with a transitional origin giving rise to inflammatory-effector or antigen-presenting and regulatory branches (Fig. 3d). Along pseudotime, early genes (*Arg1, Ctsb, Itgam, Lgals1, Prdx1*) reflected acute inflammation, phagocytosis, and oxidative stress; intermediate phases featured migration and remodeling (*Ly6a, Mmp8, Ace, Lyst*); later phases showed sustained inflammation with emerging immunoregulation (*Tnf, Pla2g2d, Clec4b1, Igf1, Txnip*) (Fig. 3e).

Transcription factor (TF) analysis further revealed subcluster-specific regulatory programs. M_01 (*Sox4* and *Mafb*) consistent with differentiation and metabolic regulation. M_02 (*Nuak, Etv1, Nfatc2, and Etv5*) reflected roles in homeostasis and energy pathways. M_03 (*Stat2, Irf9, Sap30, Zbtb45, and Rest*) implicated them in sustained inflammation and antiviral responses at 72 h. M_04 (*Klf13, Pbx1, Gabpb, Mxd1, and Vdr*) aligning with inflammation amplification. In contrast, M_05 (*Bach2, Batf3, Runx2, and Zeb1*) supporting antigen presentation and tolerogenic differentiation (Fig. 3c).

At the spatial transcriptomics level, we further mapped the tissue distribution of macrophage subclusters after SAH. M_01 occupied the puncture site and brain borders at 24 h, concentrating at the tract by 72 h, suggesting a close association with lesion-associated inflammation and tissue responses. M_02 remained relatively stable, consistently localized to ventricular boundaries, in line with its identity as border-associated macrophages. M_03 markedly accumulated around the puncture site at 72 h, while M_04 was broadly distributed across the parenchyma at 24 h but diminished thereafter (Fig. 3f).

### Neutrophil subcluster heterogeneity underlying immune-inflammatory programs after SAH

As core effectors of the acute inflammatory response, neutrophils exhibited striking recruitment kinetics and phenotypic transitions after SAH^16^. We captured a total of 2,355 neutrophils, with rare cells in sham brains—as expected for homeostasis—followed by a sharp rise at 24 h and a marked decline by 72 h (Fig2b). Unsupervised clustering resolved four transcriptionally and functionally distinct subsets (Fig. 4a). N_01, dominant at 24 h, expressed *Cxcl2, Ccl6, Spp1* and *Smox*, representing an acute pro-inflammatory state. N_02, predominant at 72 h, was enriched for *Apoe, Ly86, Trem2* and ribosomal genes, consistent with phagocytic and immunoregulatory functions. N_03, marked by *Camp* and *Ngp*, was relatively enriched among the rare sham neutrophils (baseline-like features), whereas N_04, characterized by *Irf7* and *Stat1*, represented an interferon-responsive subset^17^(Fig. 4a).GO enrichment delineated complementary roles: N_01 aligned with hypoxia- and injury-response pathways, consistent with its acute inflammatory profile. N_02 was enriched in phagocytosis and antiviral defense, suggesting roles in clearance and regulation during later stages. N_03 was associated with leukocyte migration and chemotaxis, reflecting early mobilization. N_04 was enriched in antiviral and endogenous inflammatory responses (Fig. 4b).

**Figure 4.**
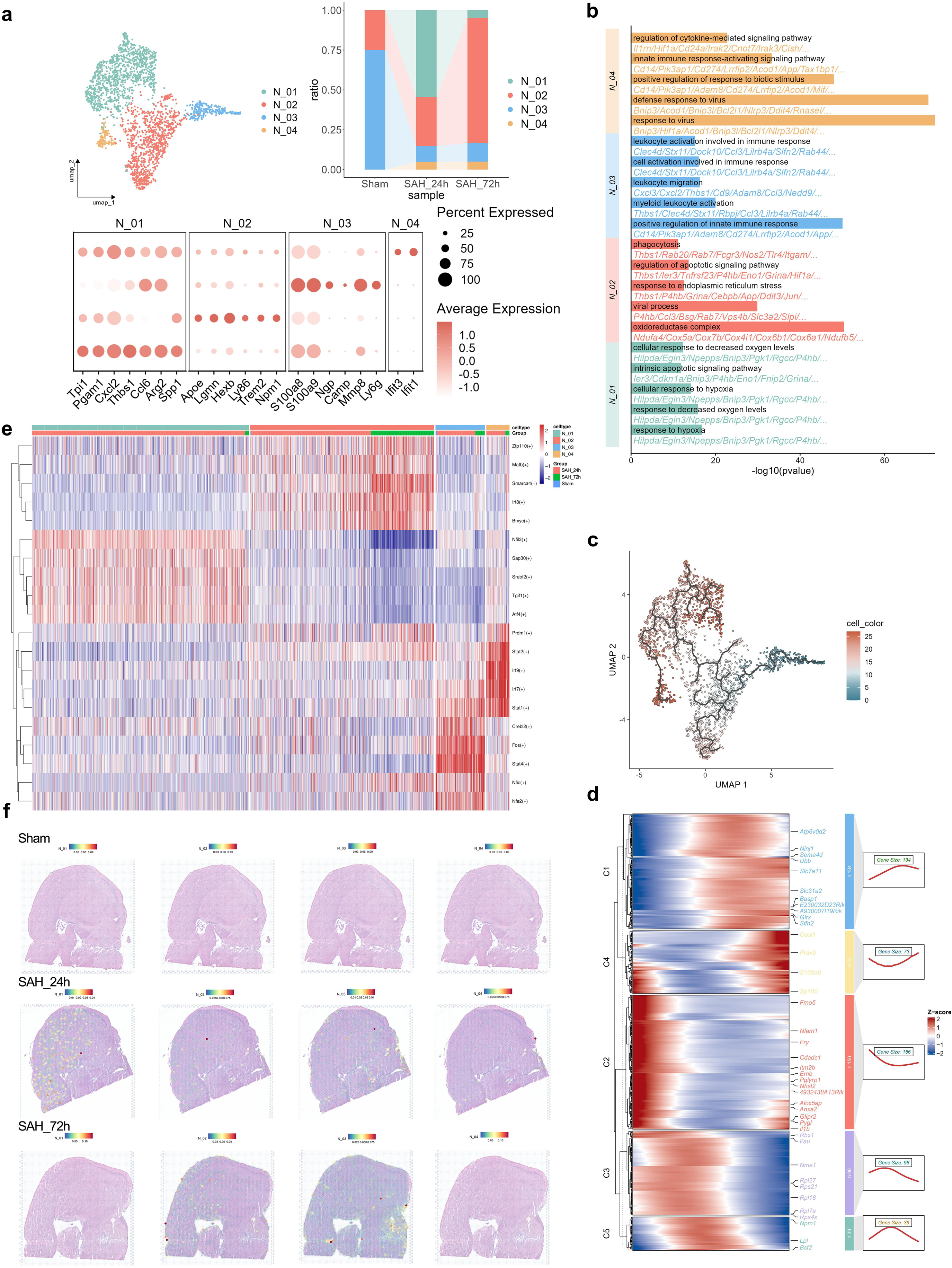
Neutrophil subcluster diversity and state transition during early SAH. a, UMAP resolving four neutrophil subclusters (N_01–N_04); Dot plot of representative markers per subcluster. d, Heatmap of the top 10 highly expressed genes within each subcluster; Subcluster abundances across conditions (sham, 24 h, 72 h). b, GO enrichment of DEGs across neutrophil subclusters (BH-adjusted *P* < 0.05). c, Pseudotime trajectories reconstructed with Monocle3. d, Heatmap of pseudotime-dependent gene dynamics (Z-scored), grouped into modules with representative genes indicated. e, Heatmap of transcription-factor activity across subclusters (top five TFs per subcluster). f, Spatial deconvolution maps of N_01–N_04 in sham, SAH_24 h, and SAH_72 h sections; color intensity reflects relative abundance.

To further dissect neutrophil development, we constructed pseudotime trajectories. Trajectory inference positioned N_03 (*Ngp⁺, Camp⁺*) near the root, diverging along two main branches: one maturing toward N_02 (phagocytic/regulatory) and the other toward N_01 (acute pro-inflammatory) or N_04 (interferon-responsive) (Fig. 4c). Along the continuum, early-stage cells expressed *Arg1, Pglyrp1, Prdx1*, (initiation/clearance), intermediate stages featured *Ly6a, Mmp8* (migration/remodeling), and late stages upregulated *Tnf, Pla2g2d, Slc7a11* (immune regulation) (Fig. 4d). Together, these results delineate a temporally ordered program that begins with acute inflammation, proceeds through migration and remodeling, and culminates in immunoregulation^18^.

Transcription factor analysis corroborated this progression. N_01 was enriched for Nfil3 and Atf4, regulators of inflammation and metabolism,; N_02 upregulated Irf8, Mafb, and Smarca4, associated with immune regulation,; N_03 expressed Fos and Stat4, characteristic of early activation,; while N_04 showed enrichment of interferon-responsive factors Stat1, Stat2, and Irf9^19^ (Fig. 4e).

Spatial transcriptomics mapped these subsets across the injured hemisphere. At 24 h, N_01 localized predominantly to the cortex and parenchyma. N_02, initially parenchymal, progressively migrated to the cortex and outer brain boundaries by 72 h. N_03 was broadly distributed in the parenchyma at early stages but concentrated near the puncture site by 72 h. In contrast, N_04, although limited in number, consistently localized to the hemorrhagic core, suggesting a focal role in antiviral and endogenous inflammatory responses (Fig. 4f).

### Temporal transcriptomic remodeling and subpopulation landscape of MG after SAH

To comprehensively delineate the dynamic responses of MG following SAH, we first profiled stage-specific transcriptomic remodeling across distinct time points. At 24 h post-SAH (vs sham), MG rapidly shifted from a homeostatic to a pro-inflammatory state^20^. This transition featured induction of chemokines and membrane-associated signaling molecules (e.g., *Ccl1, Ccl4, Plek, Nav3, Plxdc2, Rps2/Rpsa*) and repression of homeostatic and growth factor receptor–like genes (*Csf1r, P2ry12, Selplg*), together with immediate-early/stress markers (*Jun, Rhob, Btg2, Sparc, Ttr*) (Fig. 5a). Pathway analysis indicated activation of TNF, NF-κB, and Toll-like receptor (TLR) signaling, cytosolic DNA sensing, HIF-1 signaling, chemotaxis/endocytosis, and apoptosis, accompanied by broad suppression of multiple metabolic axes (fatty-acid metabolism, peroxisome, oxidative phosphorylation, TCA/pyruvate metabolism and lipid programs), consistent with early immuno-metabolic reprogramming (Fig. 5b).

**Figure 5.**
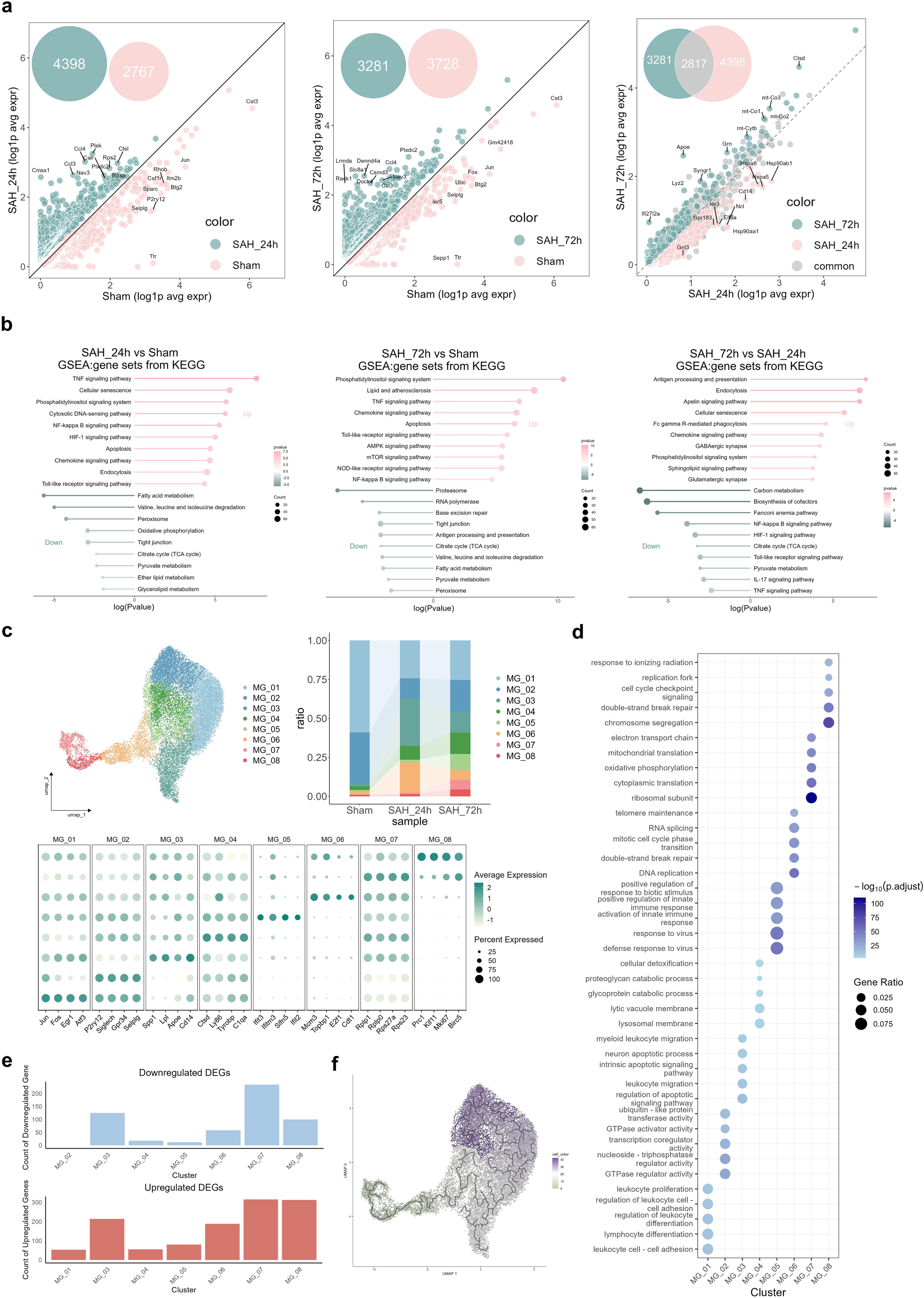
Microglial heterogeneity and functional states in early brain injury after SAH. a, Volcano plots of microglial DEGs for SAH_24 h vs sham, SAH_72 h vs sham, and SAH_72 h vs SAH_24 h, with Venn/Euler insets summarizing overlap of upregulated genes across comparisons (Wilcoxon, two-sided; BH-adjusted *P* < 0.05,log₂FC| > 0.25, min.pct ≥ 0.10). b, KEGG pathway enrichment of DEGs from the three contrasts (FDR < 0.05). c, UMAP resolving eight transcriptionally distinct microglial subclusters (MG_01-MG_08); Subcluster abundances across conditions (sham, 24 h, 72 h); Dot plot of representative markers per subcluster. d, GO enrichment of microglial subcluster-specific DEGs (BH-adjusted *P* < 0.05). e, Bar plots showing the numbers of upregulated and downregulated genes per microglial subcluster. f, Pseudotime trajectories reconstructed with Monocle3.

By 72 h post-SAH (vs sham), MG maintained cytoskeletal/endocytic remodeling (*Rack1, Dock4, Dennd4a, Slc8a1*) and chemokine activity (*Ccl3, Ccl4*), while homeostatic and nutrient-transport markers (*Selplg, Sepp1, Ttr*; with reduced *Fos, Jun, Btg2, Ubc*) remained low, indicating persistent inflammatory stress. Direct comparison 72 h vs 24 h suggested a partial shift toward clearance/repair: lipid metabolism, lysosomal and mitochondrial programs (upregulation of *Apoe, Ctsd*, *mt-Co1/2/3*, *mt-Cytb* and *Lyz2*), alongside synaptic regulators (*Syngr1*), together with downregulation of stress chaperones and pattern-recognition components (*Hspa8, Hspa5, Cd14*) and nucleolar/chromatin factors (*Ncl, Gpr183, Gnl3*). Concordantly, while TNF/NF-κB/TLR signaling remained active at 72 h, additional enrichment emerged in AMPK–mTOR and NOD-like receptor pathways, underscoring strengthened links between metabolic sensing and effector immunity. Relative to 24 h, antigen processing/presentation, endocytosis, and Fcγ receptor–mediated phagocytosis were markedly enhanced, accompanied by increased phosphatidylinositol/sphingolipid signaling and neurotransmission-related programs (GABAergic and glutamatergic synapses) (Fig. 5a,b).

Within this overall framework, unsupervised clustering on the 20,281 MG resolved eight subpopulations with distinct phenotypes: early-response MG (MG_01; *Jun^+^, Fos^+^*), homeostatic MG (MG_02; *P2ry12^+^, Selpl^+^*),, inflammatory MG (MG_03; *Spp1^+^, Lpl^+^*), complement-associated MG (MG_04; *C1qa^+^, Ctsd^+^*), interferon-associated MG (MG_05; *Ifit3^+^, Ifit2^+^*), proliferative MG1 (MG_06; *Mcm3^+^, Topbp1^+^*), proliferative MG2 (MG_07; *Rplp1^+^, Birc5^+^*), and proliferative MG3 (MG_08; *Mki67^+^, Birc5^+^*)(Fig. 5c). GO enrichment analysis revealed that MG_01 was enriched in processes related to leukocyte adhesion, differentiation, and proliferation; MG_02 was closely associated with GTPase regulation, transcriptional co-regulation, and ubiquitin-like protein transfer; MG_03 mainly involved apoptotic signaling regulation, myeloid leukocyte migration, and neuronal apoptosis; MG_04 participated in lysosomal and vacuolar membrane processes, glycoprotein and proteoglycan catabolism, as well as cellular detoxification; MG_05 was strongly linked to antiviral defense and innate immune activation; MG_06 was enriched for DNA replication, double-strand break repair, mitotic cell cycle transition, RNA splicing, and telomere maintenance; MG_07 was associated with ribosomal subunits, cytoplasmic and mitochondrial translation, oxidative phosphorylation, and the electron transport chain; while MG_08 mainly participated in chromosome segregation, double-strand break repair, cell-cycle checkpoint signaling, replication fork maintenance, and responses to ionizing radiation(Fig. 5d).

Collectively, these data depict a temporal sequence in which MG rapidly abandon homeostasis for inflammatory programs at 24 h, then partially reprogram toward lipid handling, lysosomal–mitochondrial coupling and phagocytic/antigen-presentation functions by 72 h, while a set of proliferative states (MG_06–MG_08) expands, indicating coordinated inflammatory and biosynthetic remodeling that has yet to fully normalize.

### From molecules to space: functional and lineage remodeling of MG subpopulations

Based on upregulated and downregulated DEGs, MG_03, MG_06, MG_07, and MG_08 exhibited the highest activation levels. Together with prior reports, these four subpopulation were designated as SAH-associated microglial subpopulations (SAMs)^21,22^ (Fig. 5e).

Pseudotime analysis revealed a well-defined differentiation trajectory of MG. MG_08 resides near the root, giving rise to two primary branches toward MG_07 and MG_06. From MG_06, the trajectory bifurcated again: one branch progressed toward MG_05/MG_04, representing clearance- and complement/adhesion-related states, while the other directed toward MG_03, which subsequently differentiated into MG_01 (early-response) and MG_02 (homeostatic)(Fig. 5f). Along this continuum, early-stage cells were enriched for proliferative and spindle assembly genes (*Birc5, Mcm7, Mad2l1, Tpx2*), intermediate stages for response and antigen-presentation markers (*Anxa2, Il1a, Cd83*), and late-stage cells were characterized by clearance and immunomodulatory programs (*Emp1, Nlrp3, Serinc3*), outlining a continuous transition from proliferation to response amplification and ultimately to clearance with immunomodulatory programs (Fig. 6a).

**Figure 6.**
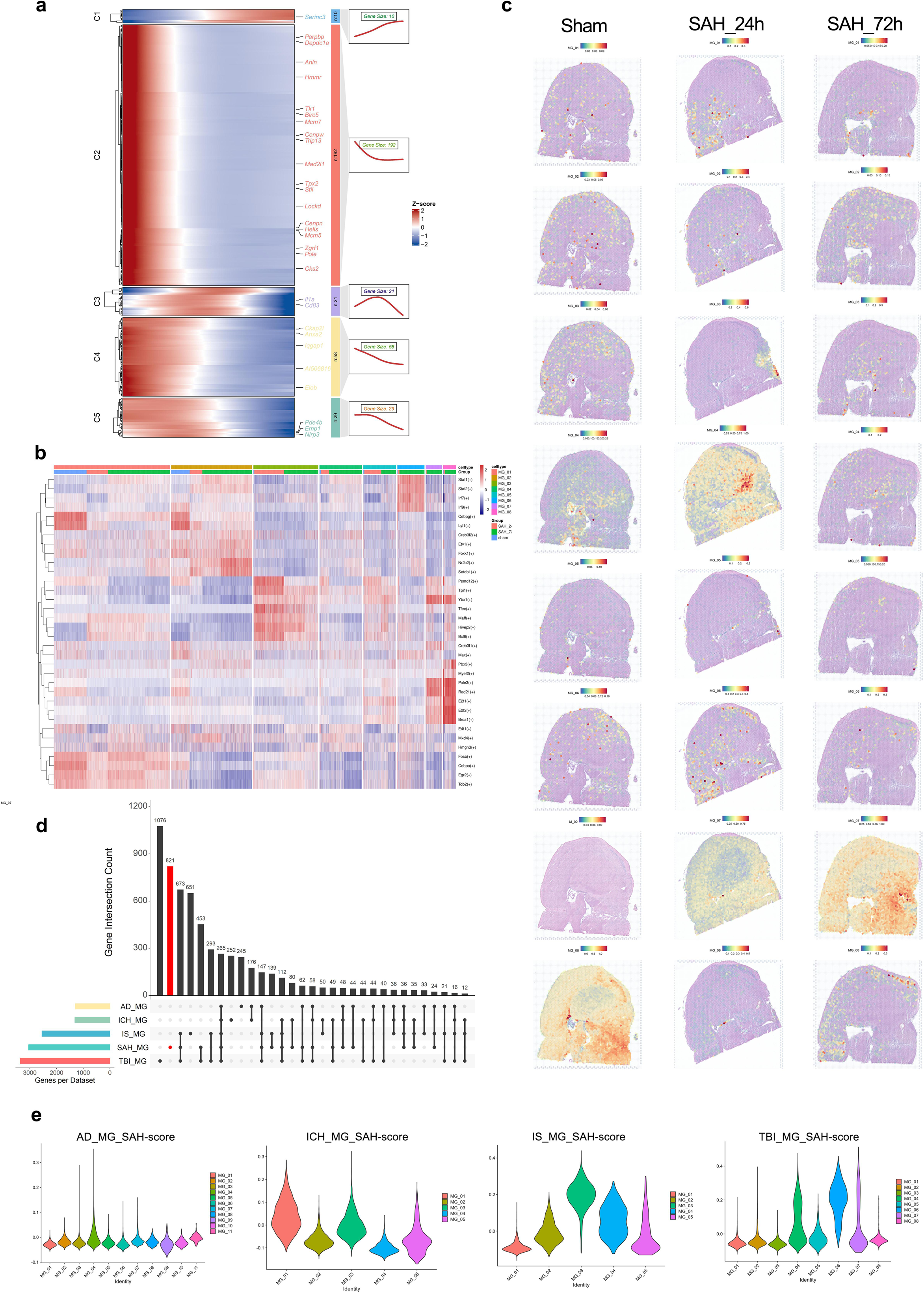
Integrated transcriptomic and regulatory landscape of microglial subclusters and the emergence of SAH-specific MG_03. a, Heatmap of pseudotime-dependent gene dynamics (Z-scored), grouped into modules with representative genes indicated. k, Heatmap of regulon/TF activity across microglial subclusters (pySCENIC/AUCell), displaying top enriched transcription factors per subcluster. c, Spatial deconvolution maps of microglial subclusters (MG_01–MG_08) across sham, SAH_24 h, and SAH_72 h sections. Color intensity reflects relative abundance per spot. d, Venn diagram showing the overlap and uniqueness of differentially expressed genes (DEGs) in SAH-specific injury-associated microglia (MG_03) compared with microglia associated with Alzheimer’s disease (AD), traumatic brain injury (TBI), ischemic stroke (IS), and intracerebral hemorrhage (ICH). e, Violin plots showing disease-associated microglial gene expression in intracerebral hemorrhage (ICH), ischemic stroke (IS), traumatic brain injury (TBI), and Alzheimer’s disease (AD).

Consistent with these stage-specific dynamics, transcription factor expression further highlighted subtype-specific regulatory features. MG_03 was marked by high expression of Tfec, Bcl6 and Ybx1, suggestive of enhanced metabolic adaptation and immune-modulatory potential. MG_06 showed elevated Stat1, Stat2 and Irf7, aligning with an interferon-driven proliferative and repair response. MG_07 displayed increased E2f1 together with Ybx1 and Rad21, not only supporting its biosynthetic and antigen-presenting functions but also suggesting a moderate proliferative potential that distinguishes it as an intermediate state between clearance-oriented and highly proliferative subsets. In contrast, MG_08 was enriched for E2f1, E2f2 and Brca1, reflecting strong mitotic activity, chromatin remodeling and genome-stability programs that positioned it at the proliferative/initiating pole (Fig. 6b).

ST further resolved distinct anatomical niches MG subtypes across time (Fig. 6c). MG_01 remained relatively stable, primarily localized to the parenchyma adjacent to the ventricles. MG_02 was mainly localized to cortex and parenchyma. MG_03 underwent the most pronounced redistribution- present diffusely in parenchyma at sham conditions, clustered at the puncture tract at 24 h, and shifted toward periventricular regions by 72 h. MG_04 was enriched near the puncture site at 24 h and expanded into broader parenchymal at 72 h. MG_05, although relatively sparse, localized around the ventricles and cortical areas. MG_06 remained parenchyma-enriched with minimal spatial drift. MG_07 was consistently tract-proximal, whereas MG_08 moved from periventricular regions at 24 h to subcortical parenchyma at 72 h. These patterns couple functional programs and lineage progression to spatial niches in early SAH.

### MG_03 defines a SAH-specific inflammatory microglial state in the acute phase

To determine whether MG_03 represents a disease-associated microglial state distinct from other central nervous system disorders, we analyzed publicly available datasets from Alzheimer’s disease (AD)^23^, ischemic stroke (IS)^24^, traumatic brain injury (TBI)^25^, and intracerebral hemorrhage (ICH)^26^, and first identified representative disease-associated microglial populations in each condition (AD_MG_04, ICH_MG_03, IS_MG_03, and IS_MG_04). We then compared differentially expressed genes (DEGs) of MG_03 in SAH with those of these disease-associated microglial subclusters (Fig. 6d). Analyses focused on the acute phase (24 h), when MG_03 was most abundant and exhibited the strongest transcriptional remodeling in SAH (Fig. 5c,d).

Analysis revealed that MG_03 harbors a distinct set of 794 SAH-enriched genes (Supplementary Table1) (Fig. 6d). These SAH-specific transcriptional features were primarily associated with immune sensing, inflammatory receptor signaling, and injury-responsive pathways, including damage-associated molecular pattern recognition mediated by *Tlr8*^27^, enhanced TNF receptor signaling involving *Tnfrsf9* (*Cd137*)^28^ and *Tnfrsf8* (*Cd30*)^29^, and enrichment of myeloid immune activation receptors such as *Cd300e*^30^. Together, these features delineate a highly inflammatory transcriptional state of MG_03 during acute SAH. In parallel, MG_03 shared a subset of inflammatory and stress-related genes with disease-associated microglia across conditions, constituting a conserved core program common to disease-associated microglial states (Fig. 6d).

To quantitatively assess disease specificity, we constructed a gene set based on the top 200 high-confidence MG_03 DEGs and applied AddModuleScore analysis across microglial populations from the different disease datasets (Fig. 6e). This analysis supported the validity of the selected disease-associated microglial populations and revealed distinct scoring patterns across diseases, further supporting MG_03 as a SAH-specific inflammatory microglial state during the acute phase.

We next investigated intercellular communication to characterize MG_03 interactions within the acute SAH microenvironment. Pathway-level analysis showed significant enrichment of inflammation-and immune-regulatory signaling pathways in MG_03, including CCL^31^, TNF,^32^ CADM, SPP1, CSF, and TWEAK^33^, collectively mediating immune cell recruitment, inflammatory amplification, cell adhesion, and myeloid regulation(Fig. 7a).

**Figure 7.**
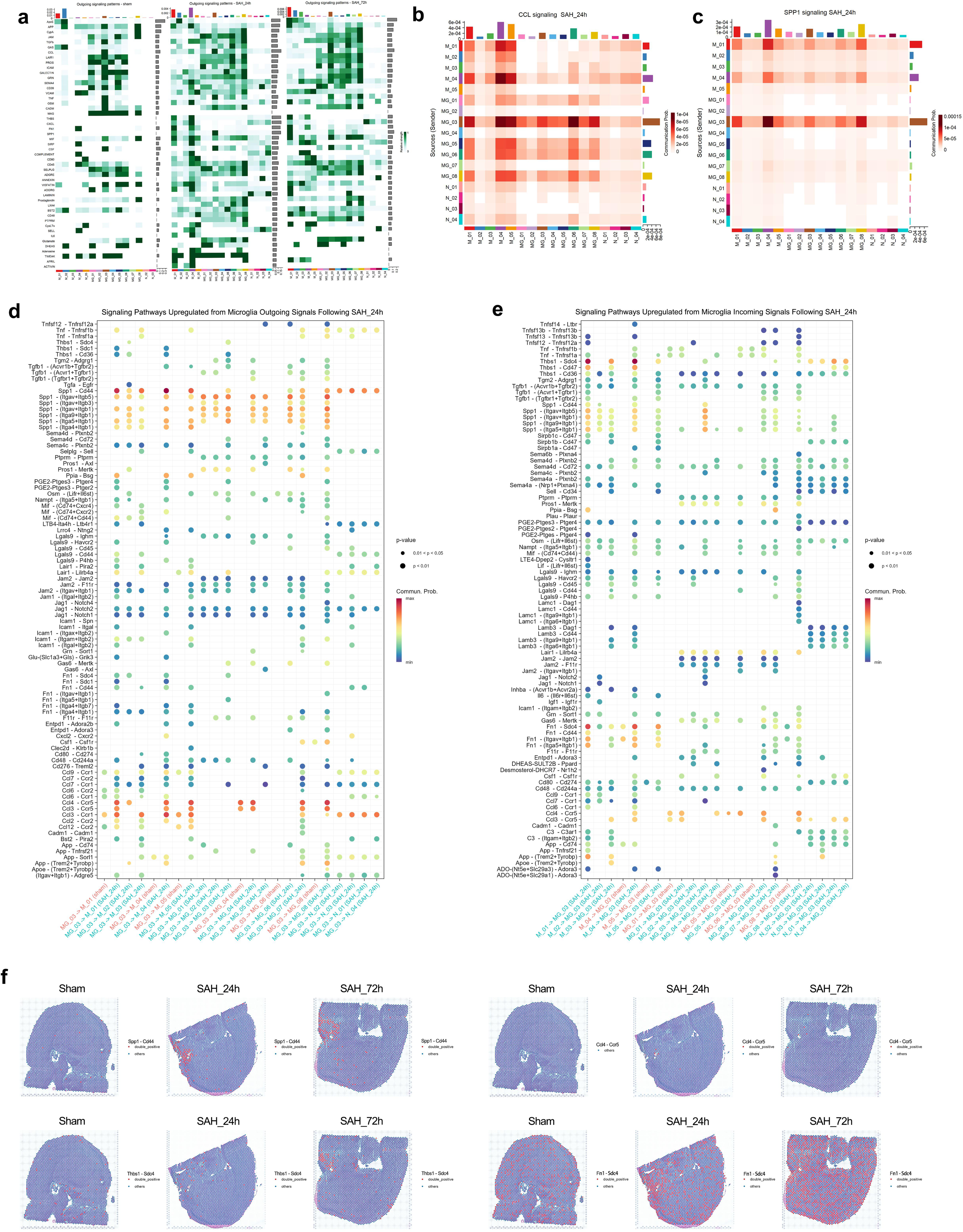
Subcluster-resolved intercellular communication landscape in SAH. a, Heatmap illustrating outgoing signaling patterns of macrophage, microglial, and neutrophil subpopulations.b, Heatmap depicting the expression levels of genes involved in the CCL signaling pathway across microglial subclusters. c, Heatmap showing the expression of genes associated with the SPP1 signaling pathway across microglial subclusters. d, Dot plot illustrating ligand–receptor interactions originating from MG_03 toward other cell populations. Dot size indicates the interaction probability, and color intensity reflects the mean expression level of the ligand–receptor pair. e, Dot plot showing ligand–receptor interactions targeting MG_03 from other cell populations, with dot size representing interaction probability and color intensity indicating mean expression. f, Spatial enrichment analysis of the Spp1–Cd44 ligand–receptor pair. g, Spatial enrichment analysis of the Ccl4–Ccr5 ligand–receptor pair. h, Spatial enrichment analysis of the Thbs1–Sdc4 ligand–receptor pair. i, Spatial enrichment analysis of the Fn1–Sdc4 ligand–receptor pair.

Ligand–receptor analysis indicated that SPP1-associated interactions constituted the dominant outgoing signaling module of MG_03. Specifically, Spp1–Cd44^34^ and multiple Spp1–integrin pairs were highly enriched in interactions between MG_03 and inflammatory macrophage subsets, particularly M_01, M_03, and M_04, implicating adhesion, migration, and inflammation-associated tissue remodeling(Fig. 7c,d). In addition, CCL–CCR signaling^31^ (e.g., Ccl2–Ccr2 and Ccl3/Ccl4–Ccr5) represented another major outgoing axis, preferentially targeting M_01 and M_05 subsets and reflecting chemokine-mediated immune cell recruitment(Fig. 7b,d).Incoming signals to MG_03 were predominantly derived from inflammatory macrophages and were enriched for extracellular matrix– and adhesion-related interactions. Thbs1–Sdc4 interactions^35^ mainly originated from M_01, M_03, and M_04 subsets, whereas Fn1–Sdc4 signals^36^ were primarily contributed by M_01, M_04, and M_05 subsets, suggesting a role for matrix-associated cues in maintaining MG_03 activation(Fig. 7e).

To validate these inferred interactions spatially, we mapped key ligand–receptor pairs onto spatial transcriptomic data. Spp1–Cd44 exhibited robust co-expression and was highly concentrated around the puncture tract at 24 h, coinciding with MG_03 enrichment. In contrast, Ccl4–Ccr5 showed limited spatial co-localization. Among incoming interactions, Thbs1–Sdc4 displayed a diffuse perilesional distribution, whereas Fn1–Sdc4 was detected both at the puncture tract and broadly within the parenchyma (Fig. 7f).

## Discussion

In this study, we combined scRNA-seq and ST to construct a spatiotemporal immune atlas of SAH and show that immune cell states undergo a coordinated transition from acute activation to regulatory and reparative programs. Within 24 h, macrophages, neutrophils, and MG rapidly adopted pro-inflammatory phenotypes that amplify tissue injury; by 72 h, these populations progressively shifted toward antigen presentation, metabolic remodeling, and proliferative renewal. Such dynamics mirror immune responses described in other CNS disorders where an initial wave of inflammation is followed by the emergence of regulatory or repair-oriented subsets. Situating SAH within this broader framework underscores a conserved activation-to-regulation principle across neuropathologies while also highlighting features specific to hemorrhagic injury, thereby informing therapeutic strategies that exploit time-dependent windows.

Macrophages exhibited marked temporal heterogeneity, shifting from pro-inflammatory activation at 24 h to more regulatory and antigen-presenting states at 72 h. The acute-phase subset M_04 (*Cxcl2⁺, Il1b⁺*) was broadly distributed and likely drove early injury via chemotaxis and cytokine release, whereas M_03 (*Ifit3⁺, Irf7⁺*) and M_05 (*H2-Aa⁺, H2-Ab1⁺*) predominant by 72 h, enriched for lysosomal and MHC-II pathways. Similar trajectories have been reported in ischemic stroke, where infiltrating macrophages initially express chemokines (e.g., *Ccl2,Cxcl2*) before transitioning toward antigen presentation and repair-associated programs^37^. In intracerebral hemorrhage, osteopontin-expressing macrophages resembling our *Spp1⁺* transitional subset (M_01) have been linked to tissue remodeling. DAM-like populations in Alzheimer’s disease express *Spp1*, *Lgals3*, and lipid metabolism genes, closely paralleling subsets observed here^38^. Notably, our spatial add a lesion-context dimension: M_04 was parenchyma-wide early, whereas M_03 later clustered at the puncture tract-suggesting opportunities to redirect macrophage fates at defined windows.

Among the infiltrating immune cells, neutrophils stood out for their rapid yet transient response: they were most abundant at 24 h but largely declined by 72 h, consistent with short-lived but potent role in acute inflammation. The N_01 subset (*Cxcl2⁺, Thbs1⁺, Lcn2⁺*) dominated early with strong pro-inflammatory and chemotactic signatures, aligning with rapid tissue infiltration, degranulation, and amplification of the inflammatory cascade; analogous populations have been linked to blood–brain barrier disruption and neuronal apoptosis in ischemic stroke^39^, and to vascular damage in intracerebral hemorrhage^40^. The interferon-responsive subset N_04 (*Irf7⁺, Stat1⁺*), although smaller, localized to the hemorrhagic core and resembled IFN-driven neutrophils described in viral encephalitis and Alzheimer’s disease, where they contribute to antiviral defense and inflammatory amplification. By 72 h, N_02 acquired phagocytic/immunoregulatory features reminiscent of N2-like neutrophils reported in gliomas, which can modulate immunity and tissue remodeling^41^. These parallels support a model in which neutrophils act as “first responders”, and in some contexts, may adopt regulatory states that shape later outcomes. Targeting these early inflammatory states without impairing their potential regulatory transition could represent a critical therapeutic strategy in SAH and related pathologies.

MG, the resident CNS macrophage, underwent pronounced transcriptional and spatial remodeling in SAH, transition from homeostatic and early-response states to specialized inflammatory and proliferative programs. We identified four SAMs populations: MG_03 (*Spp1⁺, Lpl⁺*), enriched at 24 h, represented a strongly inflammatory state linked to early tissue injury, whereas MG_06, MG_07, and MG_08 expanded by 72 h with cell-cycle and biosynthetic signatures. Spatial mapping further revealed distinct niches: MG_03 concentrated at the puncture site acutely, while proliferative subsets redistributed to periventricular and subcortical regions later. These features echo microglial phenotypes described in other neurological disorders. DAMs in Alzheimer’s disease share *Spp1*, *Lpl,* and lipid metabolism signatures with MG_03 that drive neuroinflammation^42^, and late DAM subpopulations enriched for cell-cycle and biosynthetic pathways modules paralleling MG_06–08. In multiple sclerosis, lesion microglia show elevated MHC-II, parallelling to antigen-presenting capacity of observed here (notably within MG_07)^43^. TBI induces proliferative microglia resembling MG_06 and MG_08, which replenish depleted microglial pools and support repair, and gliomas harbor metabolically heightened microglia with oxidative phosphorylation and translation signatures, akin to MG_07. Thus, inflammatory and proliferative axes of microglial specialization appear conserved, whereas the SAH-specific spatial redistribution—early clustering of inflammatory SAMs at the lesion site, followed by broader dispersion of proliferative SAMs—may be characteristic of hemorrhagic injury. This raises the possibility that selectively suppressing MG_03 while fostering proliferative/clearance-competent states could tilt the balance toward repair. Notably, cross–neurological disease comparative analyses further indicate that the transcriptional profile of MG_03 is more tightly coupled to acute injury sensing and inflammatory amplification processes, thereby distinguishing it from the more broadly defined disease-associated microglial states described in other conditions^24^. In line with this interpretation, intercellular communication analyses position MG_03 as a localized inflammatory signaling hub within the early SAH microenvironment, where it preferentially engages inflammatory macrophage subsets through adhesion-related, extracellular matrix–associated, and chemokine-mediated signaling pathways.

Several limitations of our study should merit consideration. First, ST profiling included limited biological replicates, which may underrepresent inter-individual variability. Second, our time frame (24 h-72 h), captures early injury but not later phases (e.g., delayed cerebral ischemia). Additionally, this study exclusively utilized male mice to minimize the potential confounding effects of the estrous cycle on acute immune responses; however, considering sexual dimorphism in SAH outcomes, future validation in female models is warranted. Finally, transcriptomic data alone cannot fully resolve functional causality. Future studies incorporating protein-level validation, longitudinal sampling, and human specimens and targeted perturbations of nominated axes will be essential to refine mechanism and clinical relevance. Expanding temporal resolution and integrating additional omics layers should further mature this atlas into a clinical translational framework.

In summary, our data delineate a coordinated trajectory in which innate immune cell populations in SAH progress from acute inflammatory activation to regulatory and reparative states. By linking distinct macrophage, neutrophil, and microglial subsets to spatial niches and lineage dynamics, this work situates SAH within the broader framework of CNS immune remodeling observed in stroke, hemorrhage, neurodegeneration, and trauma, and highlights time and targetable opportunities to attenuate early injury while harnessing later reparative programs.

## Methods

### Ethical approval

All animal procedures were approved by the second affiliated hospital of Zhejiang university - Animal Ethics Committee (Approval no. AIRB-2024-0285) and were performed according to the Guiding Principles for the Care and Use of Laboratory Animals Approved by Animal Regulations of National Science and Technology Committee of China.

### Animals

Adult male C57BL/6J mice (8–10 weeks old, body weight 22–26 g) were purchased from Shanghai SLAC Laboratory Animal Co., Ltd. Mice were housed in a specific pathogen-free facility under controlled temperature (22 ± 2 °C) and humidity (50 ± 10%) with a 12/12-h light/dark cycle. Food and water were provided ad libitum, and animals were allowed to acclimate for at least 2 weeks prior to experimental procedures. Animals were randomly assigned to experimental groups, and investigators were blinded during outcome assessment and data analysis to minimize bias.

### scRNA-seq — library preparation and sequencing

Single-cell barcoding and library construction were performed using the Chromium Single Cell 3’ v2 platform (10x Genomics). Libraries were sequenced on an Illumina NovaSeq 6000 system, targeting a depth of ≥100,000 reads per cell using a paired-end 120 bp configuration.

### ST— library preparation and sequencing

Brains were cryosectioned at 10 µm and mounted onto Visium Spatial Gene Expression slides (10x Genomics). Sections were fixed, H&E stained, and imaged prior to permeabilization. Library construction followed the Visium Spatial Gene Expression User Guide (CG000239). Final libraries were sequenced on an Illumina NovaSeq 6000 platform with a paired-end 150 bp configuration, targeting a depth of ≥100,000 reads per spot.

### DEG analysis

All differential expression analyses were performed in Seurat v5.3.0 using the two-side Wilcoxon rank-sum test, with *P* values adjusted by the Benjamini–Hochberg (BH) method. Unless noted otherwise, significance thresholds were log₂FC > 0.25, gene detected in ≥10% of cells in the tested group, and BH-adjusted *P* < 0.05. For cluster-level markers, each cluster was compared against all remaining cells (Seurat FindAllMarkers, only.pos=TRUE) and genes were ranked by effect size (log₂FC), from which representative top markers were reported. For pairwise group contrasts (e.g., sham vs SAH_24h; SAH_24h vs SAH_72h), DEGs were identified with FindMarkers under the same thresholds, retaining both up and downregulated genes; the resulting gene sets were used for downstream functional enrichment.

### scRNA-seq data preprocessing and quality control

All scRNA-seq data preprocessing and quality control (QC) were performed in R v4.5.1 with Seurat package v5.3.0 ^44^. Raw FASTQ files were processed with CellRanger (10x Genomics) to generate gene–cell matrices, which were imported via Read10X and initialized with CreateSeuratObject (default thresholds min.cells = 3, min.features = 200)^45^. Per cell mitochondrial fraction was computed with PercentageFeatureSet (pattern ^mt-), and cells were filtered by:nFeature_RNA with <200 or >9,000 percent.mt >10% and abnormally low/high nCount_RNA (flagged by FeatureScatter) to exclude low-quality cells and putative doublets. After filtering, counts were normalized with NormalizeData (LogNormalize, scale. factor = 1e4), and ScaleData was used to regress unwanted variation (e.g., nCount_RNA, percent.mt). Highly variable genes were selected with FindVariableFeatures, and dimensionality reduction was performed with RunPCA based on the top variable features. QC metrics (nFeature_RNA, nCount_RNA, and percent.mt) were visualized using VlnPlot and FeatureScatter.

### Gene regulatory network (GRN) inference

Based on the single-cell gene expression matrix and TFs expression status, a co expression module between TFs and candidate target genes was constructed using the pySHENIC software package (v.0.11.2)^46^. Use RcisTarget software to identify modules with significantly enriched regulatory factor binding motifs (Motif) in target genes, and create regulatory network modules (Regulons) containing TF and directly targeted genes. “AUCell” package was used to score the activity of each group of regulators in each single cell. Subsequently, a heatmap of normalized regulon AUC was generated using R.

### Reproducibility and software

All analyses were conducted in R 4.5.1. Core single-cell and spatial workflows used Seurat v5.3.0 (with SeuratObject v5.1.0, sctransform v0.4.2, Matrix v1.7-3) and integration with harmony 1.2.3; data containers relied on SingleCellExperiment v1.30.1 and SummarizedExperiment v1.38.1. Clustering and visualization used ggplot2 v3.5.2, patchwork v1.3.1, cowplot v1.2.0, and ComplexHeatmap v2.24.1. Functional enrichment and GSEA employed clusterProfiler v4.16.0, DOSE v4.2.0, enrichplot v1.28.4, fgsea v1.34.2, KEGGREST v1.48.1, GSEABase v1.70.0 and org.Mm.eg.db v3.21.0.Trajectory inference used monocle3 v1.4.26, slingshot v2.16.0 and TrajectoryUtils v1.16.1; differentiation potency was estimated with CytoTRACE2 v1.1.0. GRN analyses followed the SCENIC framework (GENIE3 v1.30.0, RcisTarget v1.28.1, AUCell v1.30.1, SCENIC (1.3.1). Spatial deconvolution was performed with spacexr /RCTD v2.2.1.

## Data availability

The scRNA-seq data generated in this study have been deposited in the GEO database under accession codes GSE311852 and GSE311853.Additional scRNA-seq datasets analyzed in this study were retrieved from GEO database under accession codes GSE167593, GSE174574, GSE210258, and GSE269748.

## Code availability

The code was based on the official tutorials of the packages listed, no custom code was generated.

## Acknowledgements

This work was supported by grants from National Natural Science Foundation of China (82501545 to X.W.; 82471295 and 82271301 to J.Z.).

## Author contributions

C.L., B.Z., Y.L and Q.Y. contributed equally to this work. Conceptualization: X.W., J.Z., C.L.; Formal analysis: C.L., B.Z.; Writing Original Draft Preparation: C.L., B.Z., Y.L., Y.Y., J.Z.; Writing Review and Editing: all authors; Supervision: X.C.W., C.M., G.Q., F.C., Y.L; Resources: X.W., J.Z.; Data Curation: Q.Y., Y.L., X.W.

## Disclosures

The authors declare no competing interests

## Non-standard Abbreviations and Acronyms

CNS: Central Nervous System
DAM: Disease-Associated Microglia
DEG: Differentially Expressed Gene
EBI: Early Brain Injury
GO: Gene Ontology
GSEA: Gene Set Enrichment Analysis
KEGG: Kyoto Encyclopedia of Genes and Genomes
MG: Microglia
NK: Natural Killer
SAH: Subarachnoid Hemorrhage
scRNA-seq: Single-Cell RNA Sequencing
ST: Spatial Transcriptomics
TF: Transcription Factor

## Notes

### Competing Interest Statement

The authors have declared no competing interest.

